# Bacteriophage-mediated reduction of uropathogenic *E. coli* from the urogenital epithelium

**DOI:** 10.1101/2025.10.06.680661

**Authors:** Bishnu Joshi, Jacob J. Zulk, Camille Serchejian, Zainab A. Hameed, Addison B. Larson, Austen L. Terwilliger, Deepak Kumar, Indira U. Mysorekar, Robert A. Britton, Anthony W. Maresso, Kathryn A. Patras

## Abstract

Urinary tract infections (UTIs), primarily caused by uropathogenic *Escherichia coli* (UPEC), affect millions annually. UPEC gains access to the urinary tract through mucosal reservoirs including the vaginal tract. With rising antibiotic resistance and frequent recurrence, alternative non-antibiotic strategies like bacteriophage (phage) therapy are gaining attention. We explored the potential of a lytic phage, ΦHP3, as well as a phage cocktail to decolonize UPEC from the urogenital tract using *in vitro* and *in vivo* models. Phage significantly inhibited UPEC growth in both bacteriologic medium and simulated vaginal fluid. Pretreatment of human vaginal epithelial cells (VK2/E6E7) and bladder carcinoma cells (HTB-9) with phage reduced adhesion and invasion of UPEC compared with controls. Phage treatment was further able to reduce intracellular UPEC in VK2 cells. Notably, phage pretreatment did not impact phage resistant UPEC strains, indicating that phage lysis was the primary driver of phenotypes. Live confocal microscopy confirmed interaction of phage particles with UPEC and with both epithelial cell lines. *In vivo*, daily intravaginal ΦHP3 administration in humanized microbiota mice significantly reduced vaginal UPEC burden after 4 days. Treatment with a phage cocktail also reduced vaginal and cervical tissue burdens by day 7 post-treatment. UPEC dissemination was observed to uterine and kidney tissues, but burdens were not different between phage and mock-treated groups. In conclusion, we demonstrate that phage and phage cocktails can modestly reduce UPEC urogenital colonization, highlighting the potential of phage therapy as a viable treatment option for UTI prevention.

**IMPORTANCE:** Urinary tract infections (UTIs) are among the most common infections worldwide, with millions of cases each year. Due to frequent recurrence and increasing antibiotic resistance, UTIs are becoming more difficult to treat, and non-antibiotic prevention options remain limited. The bacteria responsible for UTIs, such as uropathogenic *E. coli* (UPEC), often colonize other body sites, such as the intestines or vagina, before causing infection. In this study, we investigated whether bacteriophage (phage), viruses that infect bacteria, could reduce UPEC colonization. We found that phage treatment decreased UPEC adherence to vaginal and bladder cell lines, but only modestly reduced UPEC vaginal colonization in a mouse model. These findings suggest that phages may offer a potential strategy for UTI prevention, though further research is needed to optimize their therapeutic use.

## INTRODUCTION

Urinary tract infections (UTIs), at an estimated 400 million cases per year worldwide, are one of the most common bacterial infections(1). UTIs disproportionately afflict women, with more than half of women developing at least one UTI in their lifetime(2). Uropathogenic *E. coli* (UPEC) is the predominant agent of uncomplicated and complicated UTIs across the human lifespan(2–4). Recurrent UTIs (rUTIs), defined as re-infection within 6 months, occur in about a third of cases(5) and are often caused by the same species or strain(6, 7) suggestive of a host reservoir that reseeds infection. Simultaneous gastrointestinal and bladder detection of the same UPEC strain is observed in UTI(8), rUTI(9, 10), and even asymptomatic bacteriuria (ASB)(11) suggesting gut dissemination as a precursor to UTI. However, rUTI patients show similar patterns of fecal *E. coli* relative abundance, *E. coli* blooms, and genetic lineages as controls(12, 13). Similarly, the urinary microbiota, including prevalence of *E. coli*, is not different between patients with rUTI history compared to controls(14, 15), suggesting the bladder itself is also not a consistent reservoir for UPEC. Conversely, vaginal *E. coli* prevalence is estimated to be 11-12%(16–18) but increases up to 35-40% in women with rUTI(18, 19), up to 80% in women with acute UTI(20), with increased relative abundance also observed in ASB(21). Vaginal introital or peri-urethral *E. coli* colonization precedes UTI symptom onset(22–24) and isolates are frequently the same strain identified in urine(25). Collectively, these observations support vaginal UPEC reservoirs as a critical driver of UTI and consequently, support this as an important site to target for UTI prevention(26).

UTI is the second most common indication of antibiotic prescriptions(27–30) and antibiotic resistance rates among UPEC are on the rise globally(31). Prior antibiotic usage increases the risk for UTI by 2-5 fold(32) and the risk of acquiring a subsequent multidrug-resistant UTI(33). Recurrent cycles of infection and antibiotic treatment are also associated with elevated frequency of antibiotic resistance genes in the urogenital microbiome(14). Along with antibiotic resistance, UPEC undermines antibiotic efficacy by establishing intracellular bacterial communities within the bladder(34, 35) and the vaginal epithelium(36) which are protected from antibiotic therapy and could serve as an additional source of recurrent infection(37). Recalcitrant reservoirs, antibiotic resistance, and detrimental microbiome effects of repeated antibiotic usage necessitate non-antibiotic alternatives to decolonize UPEC from mucosal and intracellular sites.

Bacteriophage (phage), viruses that infect bacteria, are known to impact *E. coli* mucosal colonization. Expansion of endogenous *E. coli* phage and concordant decrease in relative *E. coli* abundance have been observed following fecal microbiome transplant in humans(38). Similarly, diverse populations of endogenous phage populations, predicted to be active against vaginal species including *E. coli*, are associated with vaginal bacterial composition(39–41). Phage have been explored as potential therapeutic for UTIs since their discovery more than a century ago with mixed success(42). Multiple randomized trials have demonstrated the safety of phage to treat UTI(43, 44), but phage was found non-superior to placebo controls in a double-blind randomized control trial(43). While phage-mediated *E. coli* decolonization of the gut has been explored experimentally using phage(45, 46) or phage plus non-pathogenic *E. coli* competitors(47), therapeutic use of phage or phage products to control vaginal bacterial colonization has only been minimally explored and not in the context of *E. coli*(48–50).

Hypothesizing that phage can serve as a non-antibiotic alternative to limit UPEC vaginal colonization and intracellular reservoirs, we tested the impact of UPEC targeting phage and phage cocktails on UPEC adherence, invasion, and intracellular persistence in immortalized human urogenital epithelial cells. We further tested the effects of phage and phage cocktails in altering UPEC vaginal colonization and urogenital tissue dissemination in humanized microbiota (^HMb^mice)(51, 52). We found that phage and phage cocktails reduced UPEC adherence and invasion in vaginal and bladder epithelial cells and reduced intracellular populations in vaginal cells. Despite promising *in vitro* activity, vaginal phage treatment displayed muted efficacy *in vivo* when administered following UPEC vaginal colonization.

## RESULTS

### Phage inhibit UPEC growth in simulated vaginal fluid

Although phage killing of *E. coli*, including UPEC strains, is widely characterized in bacteriologic media(53–55) and host fluids including blood(56, 57), urine (53–55), synovial fluid(58), synthetic saliva(59), and cecal fluid(46), phage activity in the context of the vaginal environment has been minimally described even though phage are readily isolated from vaginal specimens(60–62). For this study, we selected ΦHP3, a well-characterized phage with broad activity against UPEC strains and demonstrated efficacy in multiple infection models(46, 55, 63). We further tested a four-phage cocktail (ΦCocktail) containing ΦHP3, two additional UPEC targeting phage, ΦE17 and ΦES19, which show superior anti-biofilm efficacy in an *in vitro* catheter-associated UTI model(55), and ΦHP3.1, a derivative of ΦHP3 that emerged in a directed evolution experiment(54). ΦHP3.1 contains two SNP mutations in spike and longtail fiber genes which likely confer its retained infectivity against parental HP3 phage-resistant *E. coli*(54), and thus it has the capacity to counter emerging resistance. This four-phage cocktail has been used to treat extended-spectrum beta-lactamase (ESBL)-producing *E. coli* UTI in a human case report(64). UPEC cystitis strain UTI89 was mixed with ΦHP3 or the ΦCocktail at a multiplicity of infection (MOI) of 1 and incubated for 12 h in LB broth or a simulated vaginal fluid (SVF) adapted from prior work(65). In LB broth, ΦHP3 and ΦCocktail significantly reduced UPEC growth as measured by optical density for the full 12 h of culture (**Fig. 1A**), with significantly reduced area under the curve (AUC) values for both phage-exposed conditions (**Fig. 1B**). In SVF, ΦHP3 significantly reduced UPEC growth for the first 7 hours while the ΦCocktail reduced for growth for the first 9 hours, with no differences in optical density at later timepoints (**Fig. 1C**). Even so, AUC values were significantly lower in both phage-exposed conditions (**Fig. 1D**), supporting lytic activity in simulated vaginal conditions.

**Figure 1.**
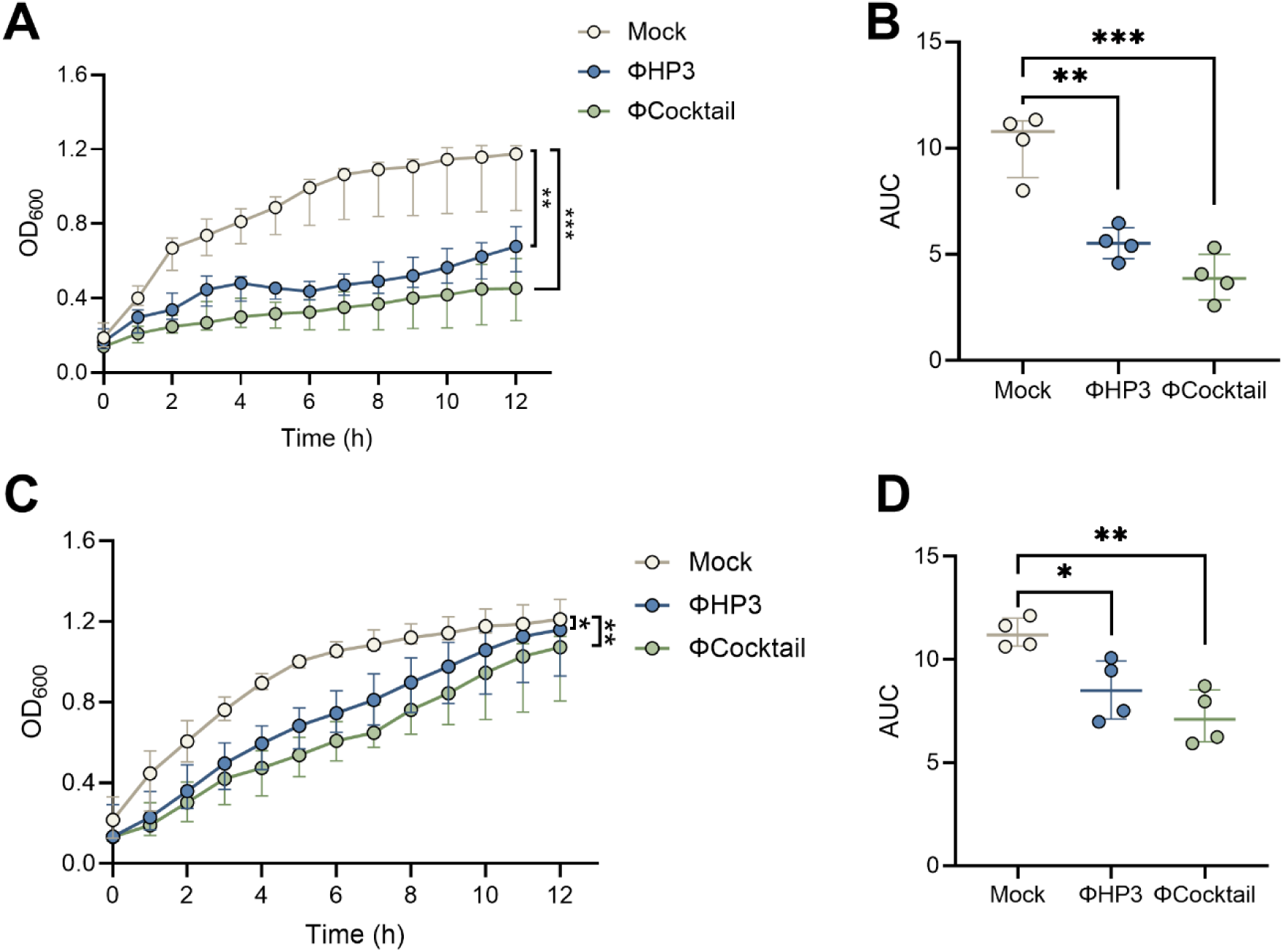
Phages inhibit UPEC growth *in vitro* in bacteriologic medium and simulated vaginal fluid. Overnight UTI89 cultures were diluted in Luria broth (LB) or simulated vaginal fluid (SVF) and infected with ΦHP3 or ΦCocktail at an MOI of 1, or mock-treated as a control. Growth was monitored by measuring OD_600nm_ at 15 min intervals for 20 h. Optical density (**A**) and area under the curve analyses (**B**) of UTI89 in LB medium. Optical density (**C**) and area under the curve analyses (**D**) of UTI89 in SVF medium. Experiments were performed in at least technical duplicate across four independent replicates. Points represent medians of four experimental replicates (A,C) or medians of individual experimental replicates (B,D). Lines represent interquartile ranges (A, C) or median with interquartile ranges (B,D). Data were analyzed by two-way ANOVA (A,C) or one-way ANOVA with Holm-Šídák’s multiple comparisons test (B,D), \**p*<0.05, \*\**p*<0.01.

### Phage decrease UPEC adherence to vaginal and bladder epithelial cells

Phage binding to eukaryotic cells, and cellular uptake of phage, varies across cell types and phage strains and may promote or inhibit phage efficacy in reducing bacterial pathogens(66–68). To assess whether phage interact with the vaginal epithelium, we performed phage adherence assays using the immortalized human vaginal epithelial cell line VK2/E6E7. ΦHP3 (10^8^ PFU) was incubated with VK2 cell monolayers for 1 hour. A subset of monolayers were washed twice with PBS to retain only cell-associated phage (Adherent Φ) whereas unwashed wells retained phage in the supernatant as well as cell-associated phage (Total Φ). Plaque assays of supernatant and cell lysates, with or without washing, revealed that washing significantly reduced cell-free PFU by 170-fold, while cell-associated PFU remained similar between conditions, at an adherence rate of 0.2% of the inoculum (**Fig. 2A**). Time lapse confocal imaging further confirmed interaction of ΦHP3, labeled with nucleic acid dye SYBR-gold(69), with VK2 cells (**Video S1**). To test the impact of phage on UPEC adherence to the vaginal epithelium, phage were incubated with VK2 cells under the conditions described in Fig. 2A and then 10^6^ CFU of UTI89 were added to the monolayers and allowed to adhere for 30 min. Both ΦHP3 and ΦCocktail pretreatment reduced UPEC adherence, 10-fold and 150-fold, respectively in unwashed conditions (Total Φ) whereas only ΦCocktail reduced UPEC adherence 40-fold when non-adherent phage were removed prior to bacterial inoculation (Adherent Φ)(**Fig. 2B**). These results suggest that both eukaryotic cell-bound and unbound phages contribute to reduced bacterial adhesion, either by directly lysing bacteria or by interfering with their ability to attach to host cells. To test whether lytic activity was required for phage-mediated UPEC reduction, we performed adherence assays using a ΦHP3-resistant UTI89 derivative (Φ^R^) with a spontaneous mutation in *rfaH* resulting in a truncated inner LPS core, the putative ΦHP3 receptor(53). No differences in UTI89 Φ^R^ adherence to VK2 cells were observed with ΦHP3 treatment in either washed (Adherent Φ) or unwashed (Total Φ) conditions (**Fig. 2C**). Confocal fluorescence microscopy further corroborated these quantitative findings. SYBR-gold labeled ΦHP3 could be visualized in close proximity with VK2 cells in the presence or absence of RFP-expressing UTI89, frequently co-localizing with bacteria in the latter condition (**Fig. 2D**).

**Figure 2.**
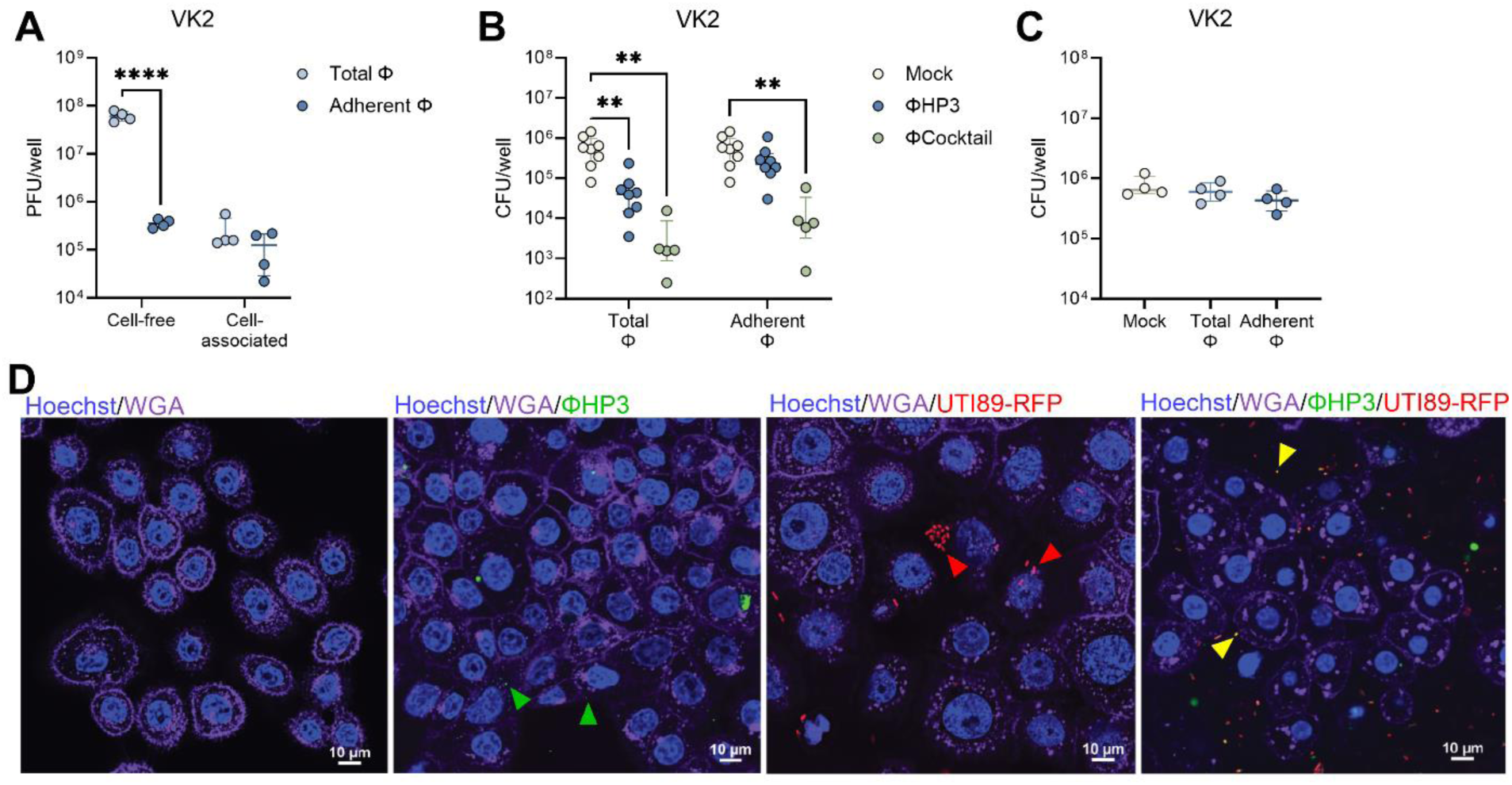
Phage pretreatment inhibits UPEC adherence to human vaginal epithelial cells. Human vaginal epithelial VK2 cell monolayers were pretreated with ΦHP3 and ΦCocktail (10^8^ PFU) for 1 hour and non-adherent phage was removed by washing (Adherent Φ) or unwashed to retain all phage (Total Φ). (**A**) ΦHP3 PFU recovered from filtered supernatant (cell-free) or lysed cells (cell-associated) for Adherent Φ and Total Φ conditions. Monolayers were infected with 10^6^ CFU of UPEC for 30 min followed by cell lysis and plating to quantify adherent bacteria. (**B**) Adherence of phage-sensitive WT UTI89 to ΦHP3 and ΦCocktail pretreated cells under Adherent Φ and Total Φ conditions. (**C**) Adherence of phage-resistant UTI89 Φ^R^ to ΦHP3 pretreated cells under Adherent Φ and Total Φ conditions. For confocal microscopy, VK2 monolayers were infected with 10^6^ CFU UTI89-RFP (red) for 3 hours, prior to treatment with 10^8^ PFU SYBR-Gold labeled ΦHP3 (green) with image acquisition immediately after phage addition. (**D**) Representative images with cell nuclei stained with Hoechst (blue) and cell membranes visualized with wheat germ agglutinin (magenta). Arrows indicate phage co-localization with host cells (green), UPEC co-localization with host cells (red), or phage co-localization with UPEC (yellow). Experiments were performed in at least technical duplicate across four to eight independent replicates (A-C) or twice independently (D). Points represent medians of experimental technical replicates and lines represent medians with interquartile ranges (A-C). Data were analyzed by two-way ANOVA with uncorrected Fisher’s LSD (A), two-way ANOVA with Holm-Šídák’s multiple comparisons test (B), or Kruskal-Wallis with Dunn’s multiple comparisons test (C), \*\**p*<0.01, \*\*\*\**p*<0.0001.

Additionally, we evaluated phage interactions with immortalized human bladder carcinoma cell line HTB-9. Identical to VK2 cell assays, ΦHP3 binding was measured in unwashed (Total Φ) and washed (Adherent Φ) conditions. Plaque assays of supernatant and cell lysates, with or without washing, revealed that washing significantly reduced cell-free PFU by 250-fold, while cell-associated PFU remained similar between conditions, at an adherence rate of 0.5% of the inoculum (**Fig. 3A**). Time lapse confocal imaging further confirmed binding of SYBR Gold-labeled phage particles to HTB-9 cells (**Video S2**). Both ΦHP3 and ΦCocktail treatment reduced UTI89 adherence 6.4-fold and 6.2-fold, respectively when cell-free phage were retained in the wells, whereas cell-associated only conditions failed to reduce UTI89 adherence to HTB-9 cells (**Fig. 3B**). As with VK2 cells, reduction in bacterial adherence was dependent on phage lytic activity as no reduction in UTI89 Φ^R^ adherence to HTB-9 cells was observed in either washed or unwashed conditions (**Fig. 3C**). ΦHP3 could be visualized in close proximity to the HTB-9 cell membrane in the presence or absence of RFP-expressing UTI89, often co-localized with bacteria when present (**Fig. 3D**).

**Figure 3.**
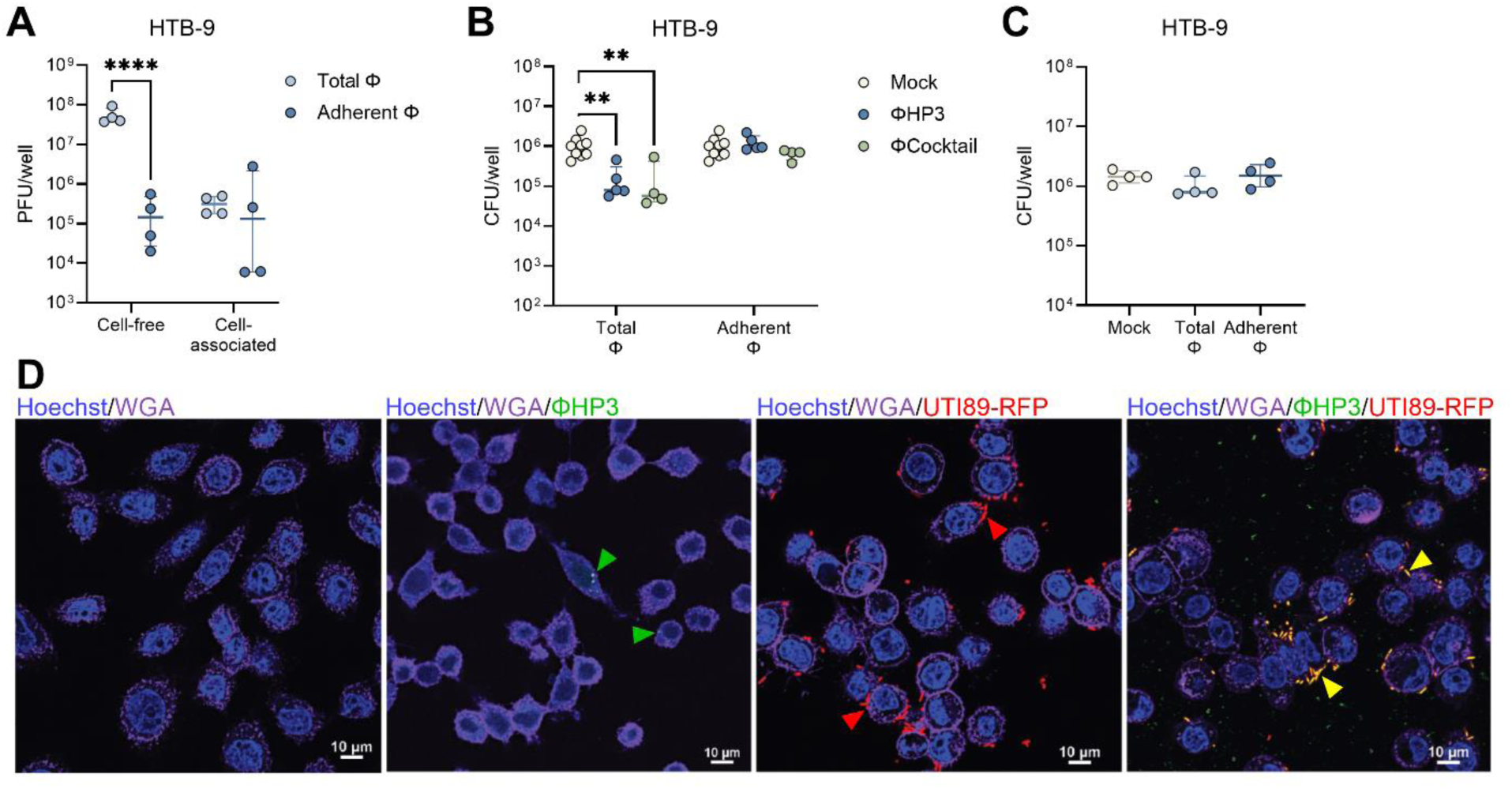
Phage pretreatment inhibits UPEC adherence to human bladder carcinoma cells. Human bladder carcinoma HTB-9 cell monolayers were pretreated with ΦHP3 and ΦCocktail (10^8^ PFU) for 1 hour and non-adherent phage was removed by washing (Adherent Φ) or unwashed to retain all phage (Total Φ). (**A**) ΦHP3 PFU recovered from filtered supernatant (cell-free) or lysed cells (cell-associated) for Adherent Φ and Total Φ conditions. Monolayers were infected with 10^6^ CFU of UPEC for 30 min followed by cell lysis and plating to quantify adherent bacteria. (**B**) Adherence of phage-sensitive WT UTI89 to ΦHP3 and ΦCocktail pretreated cells under Adherent Φ and Total Φ conditions. (**C**) Adherence of phage-resistant UTI89 Φ^R^ to ΦHP3 pretreated cells under Adherent Φ and Total Φ conditions. For confocal microscopy, HTB-9 monolayers were infected with 10^6^ CFU UTI89-RFP (red) for 3 hours, prior to treatment with 10^8^ PFU SYBR-Gold labeled ΦHP3 (green) with image acquisition immediately after phage addition. (**D**) Representative images with cell nuclei stained with Hoechst (blue) and cell membranes visualized with wheat germ agglutinin (magenta). Arrows indicate phage co-localization with host cells (green), UPEC co-localization with host cells (red), or phage co-localization with UPEC (yellow). Experiments were performed in at least technical duplicate across four to nine independent replicates (A-C) or twice independently (D). Points represent medians of experimental technical replicates and lines represent medians with interquartile ranges (A-C). Data were analyzed by two-way ANOVA with uncorrected Fisher’s LSD (A), two-way ANOVA with Holm-Šídák’s multiple comparisons test (B), or Kruskal-Wallis with Dunn’s multiple comparisons test (C), \*\**p*<0.01, \*\*\*\**p*<0.0001.

### Phage decrease UPEC invasion and intracellular survival in vaginal epithelial cells

UPEC can reside intracellularly within bladder and vaginal epithelial cells(36, 70), establishing a persistent bacterial reservoir that can serve as a nidus for urinary dissemination and recurrent infection. To determine whether phage treatment impacted UPEC invasion, we pretreated monolayers with 10^8^ PFU of ΦHP3 or ΦCocktail for 1 hour, washed away non-adherent phage (Adherent Φ) or retained non-adhered phage (Total Φ). Monolayers were then infected with UTI89 and invaded bacteria quantified using the gentamicin protection method(53). Under these conditions, intracellular bacteria could be visualized within the plasma membrane boundaries of VK2 and HTB-9 cells using confocal microscopy acquired Z-stack images after 3 h incubation (**Fig. S1**). Both ΦHP3 and ΦCocktail treatment reduced UTI89 invasion of VK2 cells 5.2-fold and 7.5-fold, respectively when cell-free phage were retained in the wells, whereas cell-associated only conditions failed to reduce UTI89 invasion (**Fig. 4A**). ΦHP3 did not alter UTI89 Φ^R^ invasion of VK2 cells (**Fig. 4B**). In HTB-9 cells, only ΦHP3 treatment significantly reduced UTI89 invasion (7.1-fold) and only when non-adherent phage were retained (**Fig. 4C**). As with VK2 cells, ΦHP3 did not alter UTI89 Φ^R^ invasion of HTB-9 cells (**Fig. 4D**).

**Figure 4.**
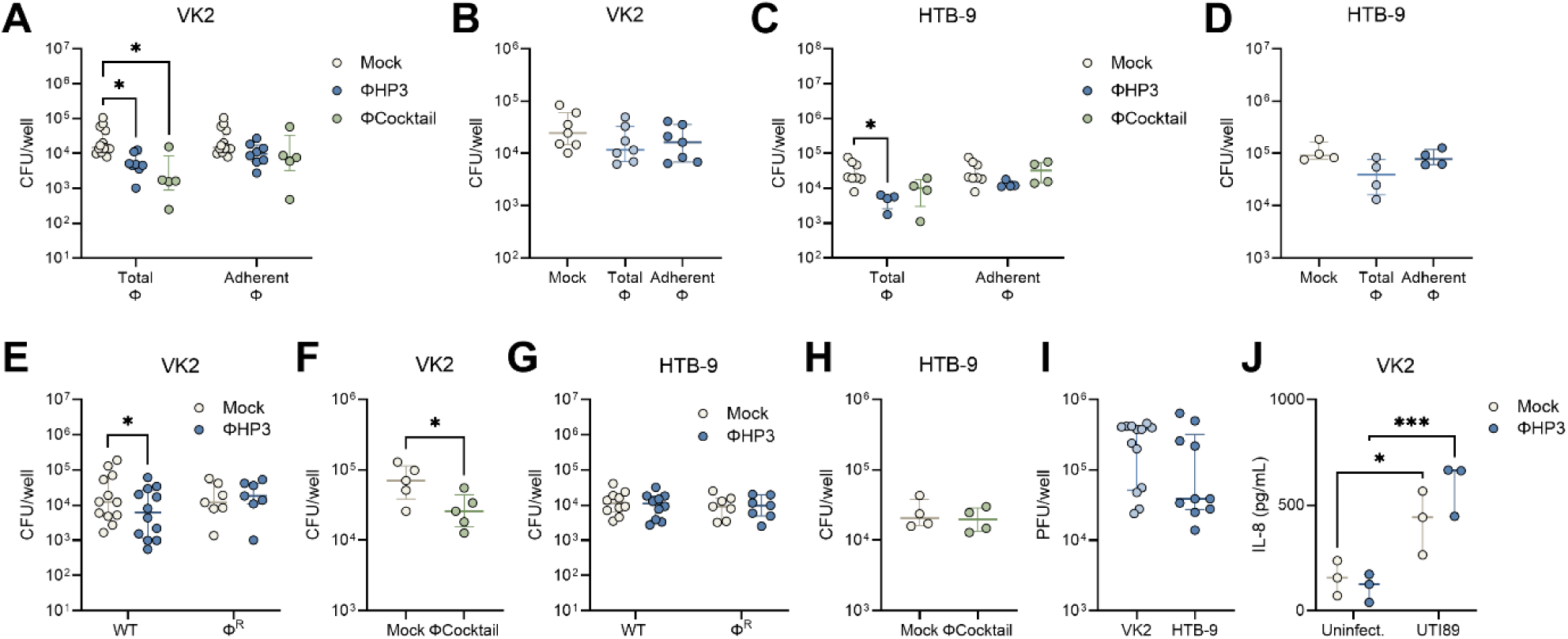
Phage modestly inhibits UPEC invasion and intracellular survival in vaginal and bladder cells. For invasion assays, human vaginal epithelial VK2 cell and bladder carcinoma HTB-9 cell monolayers were pretreated with ΦHP3 and ΦCocktail (10^8^ PFU) for 1 hour and non-adherent phage was removed by washing (Adherent Φ) or unwashed to retain all phage (Total Φ). Monolayers were infected with 10^6^ CFU of UPEC for 2 h, gentamicin treatment for 1 h to kill extracellular bacteria, followed by cell lysis and plating to quantify invasive bacteria. VK2 cell invasion of phage-sensitive WT UTI89 to ΦHP3 and ΦCocktail pretreated cells (**A**), or UTI89 Φ^R^ to ΦHP3 pretreated cells (**B**), under Adherent Φ and Total Φ conditions. HTB-9 cell invasion of phage-sensitive WT UTI89 to ΦHP3 and ΦCocktail pretreated cells (**C**), or UTI89 Φ^R^ to ΦHP3 pretreated cells (**D**), under Adherent Φ and Total Φ conditions. For intracellular survival assays, VK2 and HTB-9 monolayers were infected with 10^6^ CFU of UPEC for 2 h, gentamicin treatment for 1 h to kill extracellular bacteria. Monolayers were then treated with 10^8^ PFU phage and gentamicin for 24 h, followed by cell lysis and plating to quantify intracellular bacteria. VK2 intracellular UTI89 or UTI89 Φ^R^ after treatment with ΦHP3 (**E**), or UTI89 after treatment with ΦCocktail (**F**). HTB-9 intracellular UTI89 or UTI89 Φ^R^ after treatment with ΦHP3 (**G**), or UTI89 after treatment with ΦCocktail (**H**). (**I**) ΦHP3 adherence to VK2 and HTB-9 cells after 30 min of incubation with 10^8^ PFU, followed by washing and cell lysis. (**J**) IL-8 release in VK2 cell supernatant after 6 h of infection with 10^6^ CFU UTI89 with or without 10^8^ PFU ΦHP3 as measure by ELISA. Experiments were performed in at least technical duplicate across three to twelve independent replicates. Points represent medians of experimental technical replicates and lines represent medians with interquartile ranges (A-J). Data were analyzed by two-way ANOVA with Holm-Šídák’s multiple comparisons test (A,C,E,G), Kruskal-Wallis with Dunn’s multiple comparisons test (B,D), paired *t* test (F, H), Mann-Whitney *U* test (I), or two-way ANOVA with uncorrected Fisher’s LSD (J), \**p*<0.05, \*\*\**p*<0.001.

To test whether phage impacts established UPEC intracellular reservoirs, epithelial monolayers were infected with 10^6^ CFU for 2 hours, followed by gentamicin treatment for 1 hour, after which, 10^8^ PFU were added in fresh media containing gentamicin, and cells incubated for an additional 24 hours. Both ΦHP3 and ΦCocktail treatment modestly but significantly reduced intracellular bacteria 2.3-fold and 2.6-fold, respectively (**Fig. 4E-F**). This reduction was dependent on phage lytic activity, as UTI89 Φ^R^ intracellular survival was not impacted by phage treatment (**Fig. 4E**). In contrast, neither ΦHP3 and ΦCocktail treatment impacted UTI89 nor UTI89 Φ^R^ intracellular survival in HTB-9 cells (**Fig. 4G-H**). No differences in total phage bound to host cells within 30 minutes were observed between cell lines (**Fig. 4I**). UPEC exposure stimulates vaginal epithelial inflammation including production of chemokine IL-8(71). To test whether phage treatment altered IL-8 production in VK2 cells, monolayers were treated with 10^8^ PFU ΦHP3 for 1 hour followed by infection with 10^6^ CFU of UTI89 for 6 hours. IL-8 levels in cell supernatant were significantly increased with UTI89 exposure as measured by ELISA, but ΦHP3 pretreatment did not alter IL-8 production in infected cells or mock-infected controls (**Fig. 4J**).

### Phage treatment reduces murine UPEC vaginal colonization

To test whether phage altered the ability of UPEC to colonize the vaginal mucosa *in vivo*, we used a murine UPEC vaginal colonization model in humanized microbiota mice (^HMb^mice), generated by colonizing germ-free C57BL/6 mice with pooled human fecal microbes(52). We have previously demonstrated that ^HMb^mice harbor a more human-like vaginal microbiota that is enriched with *Lactobacillus* spp. compared to conventional mice and are readily vaginally colonized by UTI89 Strep^R^(51). As with prior studies(36, 51), mice were synchronized with β-estradiol (day -2), vaginally inoculated with UTI89 Strep^R^ (day -1), and were then vaginally administered 10^8^ PFU of phage daily for 7 days (**Fig. 5A**). Vaginal swabs were collected just prior to phage treatment and urogenital tissues were collected at day 7. By day 7, phage treatment significantly promoted UPEC vaginal clearance below the limit of detection in the ΦHP3 group (27% clearance, 10/37 mice) compared to mock-treated controls (11%, 4/37 mice, **Fig. 5B**). When comparing vaginal UPEC burdens over time, the ΦHP3 group displayed lower UPEC burdens at day 4, with no differences detected at any other time point nor in urogenital tissues at day 7 (**Fig. 5C-D**). We observed significant positive correlations between UPEC CFU across reproductive tract and kidney tissues in mock-treated mice indicating dissemination to the upper reproductive tract and urinary tract tended to co-occur within the same mice (**Fig. 5E**). However, in ΦHP3-treated mice, although positive correlations in UPEC burdens across tissues were still observed, the strength of these associations were lower and non-significant between the kidneys and the vagina or cervix (**Fig. 5F**). Since *E. coli* is detected in the ^HMb^mice vaginal microbiota(51), we tested susceptibility of endogenous vaginal *E. coli* isolates to ΦHP3 *in vitro*. All isolates tested were sensitive to phage (**Fig. 5G**). Viable plaques were recovered from most vaginal swabs over the treatment period, with levels decreasing at later timepoints (**Fig. 5H**), indicating phage retain capacity for lytic activity *in vivo*. We performed plaque assays from vaginal swabs from 10 treatment naïve mice and no plaques were observed indicating quantified PFU reflected phage treatment and not endogenous phage. Across all timepoints, vaginal UPEC CFU and ΦHP3 PFU were positively correlated suggesting that recovery of lytic phage required a viable UPEC host (**Fig. 5I**). Although *E. coli* can rapidly develop resistance to phage *in vitro* and *in vivo* (53, 54, 72, 73), all tested UPEC CFU recovered from vaginal swabs of phage-treated mice retained susceptibility to ΦHP3 and ΦCocktail (**Fig. S2A**). To test the impact of phage resistance on vaginal colonization, ^HMb^mice were vaginally colonized with UTI89 Φ^R^ (day -1) in the presence or absence of ΦHP3 treatment administered on days 0, 1, and 2. UTI89 Φ^R^ poorly colonized the vaginal tract regardless of ΦHP3 treatment, with only one mouse having detectable vaginal CFU at day 2 (**Fig. 5J**). Together, these data support that daily ΦHP3 treatment modestly reduces UTI89 vaginal colonization, driving clearance in a subset of mice over 7 days. Furthermore, persistent colonization in some animals is not due to development of phage resistance.

**Figure 5.**
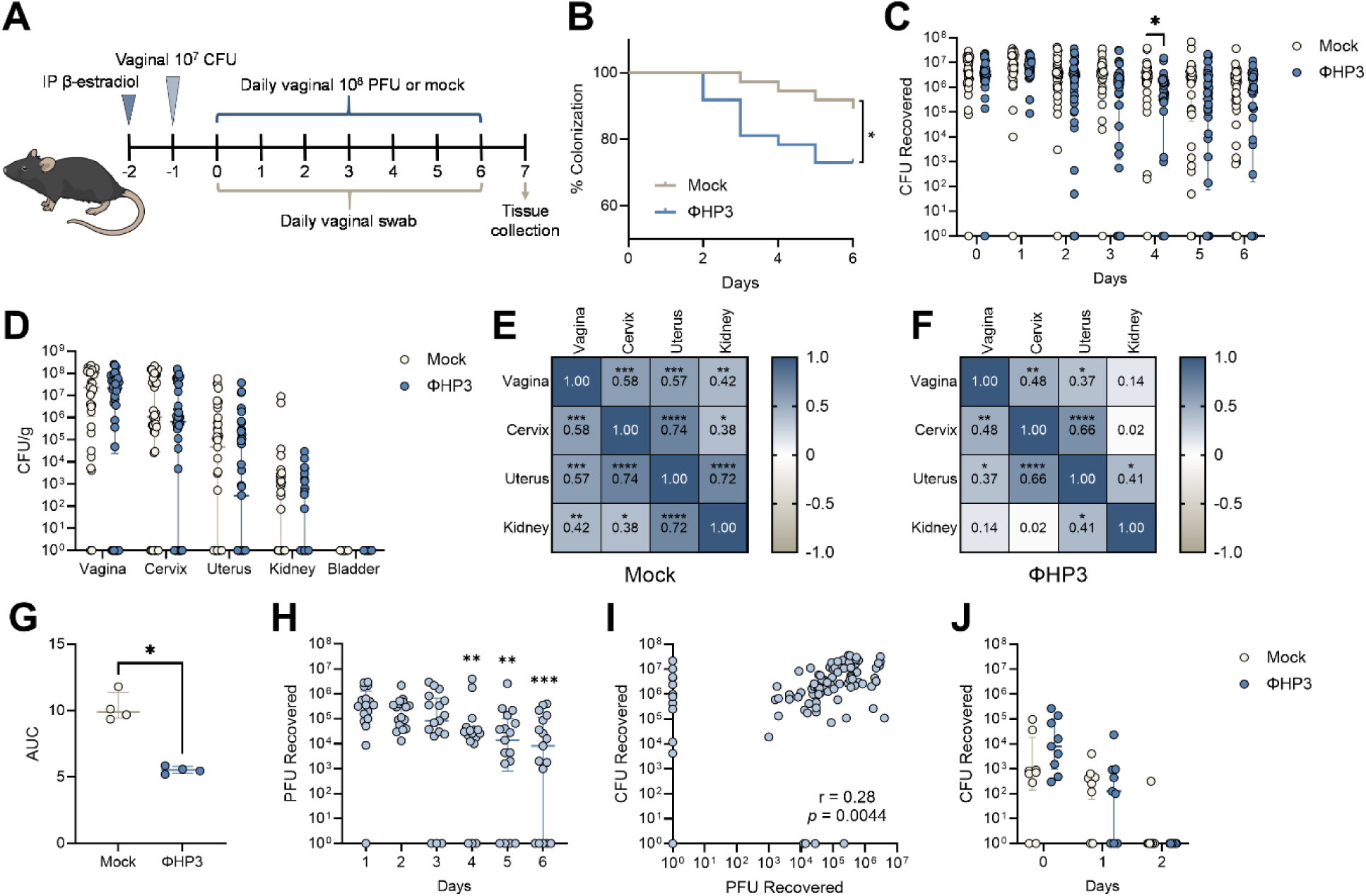
ΦHP3 reduces vaginal UPEC colonization in humanized microbiota mice. (**A**) Experimental timeline for estrus synchronization, UPEC colonization, phage treatment, and sample collection in human microbiota mice (^HMb^mice). (**B**) Percent UTI89 Strep^R^ vaginal colonization curves of ΦHP3 and mock-treatment mice over seven days of treatment, with decolonization determined as the lack of detectable UPEC on vaginal swabs for all subsequent samples. (**C**) UTI89 Strep^R^ CFU recovered from vaginal swabs over seven days of treatment. (**D**) UTI89 Strep^R^ CFU/g of urinary and reproductive tissues at day 7 post-phage treatment. Correlations of UTI89 Strep^R^ CFU across day 7 tissues in mock (**E**) and ΦHP3 treated mice (**F**). (**G**) Area under curve (AUC) of UTI89 Strep^R^ recovered from ΦHP3-treated mice vaginal swabs grown in LB for 24 h in the presence or absence of 10^8^ PFU ΦHP3. (**H**) PFU recovered from vaginal swabs on day 1 through 6 of treatment. (**I**) Correlations of PFU and UTI89 Strep^R^ CFU recovered from vaginal swabs on day 1 through 6 of treatment. (**J**) Vaginal swab CFU of UTI89 Φ^R^ Strep^R^ in the presence or absence of ΦHP3 treatment administered on days 0-2. Experiments were performed in at least two independent replicate experiments. Points represent individual mice (C, D, H-J) or medians of experimental technical replicates (G), and lines represent medians with interquartile ranges. *n*=37 (B-F), *n*=17 (H-I), *n*=9 (J). Data were analyzed by Mantel-Cox test (B), Mann-Whitney *U* test with Benjamini, Krieger and Yekutieli correction for false discovery with a false discovery rate set at 5% (C, J), two-way ANOVA with Holm-Šídák’s multiple comparisons test (D), Pearson correlation (E,F,I), Mann-Whitney *U* test (G), or Friedman test with Dunn’s multiple comparisons test to day 1 values (H), \**p*<0.05, \*\**p*<0.01, \*\*\**p*<0.001, \*\*\*\**p*<0.0001.

We further tested whether the four-phage cocktail altered UPEC vaginal colonization. Mice were inoculated with UTI89 and treated daily with phage as in **Fig. 5A**. Although 28% (8/29) of ΦCocktail-treated mice cleared vaginal UPEC by day 7, this was not statistically different from mock-treated mice with 8% (2/25) UPEC clearance (**Fig. 6A**). No differences were observed in vaginal UPEC CFU from day 1-6 of treatment; however, day 7 vaginal and cervical burdens which were significantly lower in ΦCocktail-treated mice (**Fig. 6B-C**). Similar to ΦHP3 kinetics, ΦCocktail vaginal PFU were reduced at later timepoints compared to day 1, and UPEC CFU and phage PFU were positively correlated (**Fig. 6D-E**). Because phage treatment showed only modest reductions in UPEC burden *in vivo*, despite potent *in vitro* activity, we tested several strategies to improve *in vivo* efficacy. Inoculation with 100-fold lower UPEC inoculum failed to further separate UPEC CFU colonization and dissemination over the 7-day treatment (**Fig. S2B-C**). Additionally, administering phage intravaginally just 30 minutes post-infection with UPEC did not further separate bacterial burdens between phage-treated groups and mock controls. In fact, this treatment regimen resulted in higher day 5 vaginal CFU and day 7 cervical burdens in ΦCocktail-treated mice compared to mock controls (**Fig. 6F-G**).

**Figure 6.**
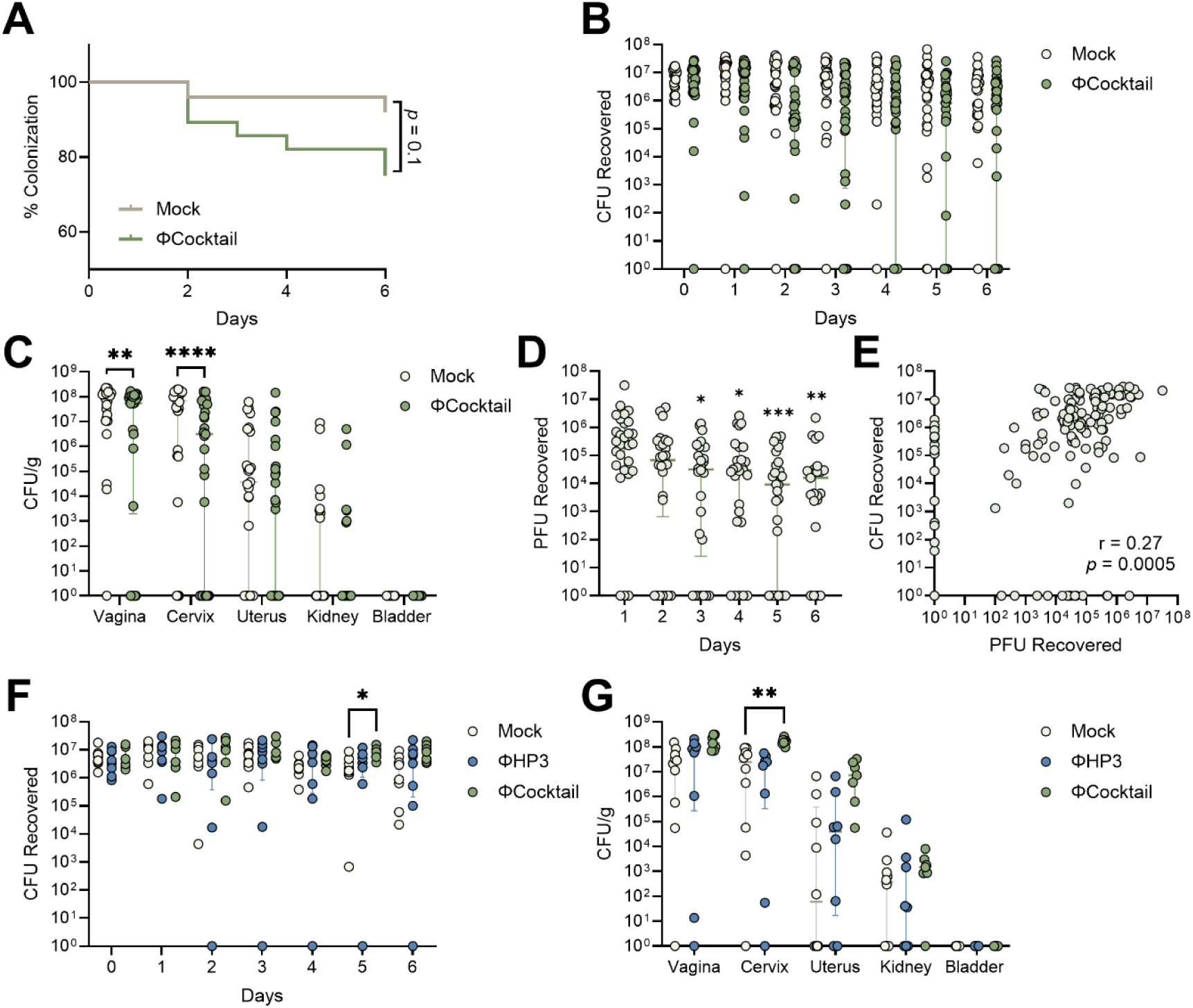
ΦCocktail modestly affects vaginal UPEC colonization in humanized microbiota mice. ^HMb^mice were vaginally inoculated with UPEC and treated daily with phage as described in Fig. 5A. (**A**) Percent UTI89 Strep^R^ vaginal colonization curves of ΦCocktail and mock-treatment mice over seven days of treatment, with decolonization determined as the lack of detectable UPEC on vaginal swabs for all subsequent samples. (**B**) UTI89 Strep^R^ CFU recovered from vaginal swabs over seven days of treatment. (**C**) UTI89 Strep^R^ CFU/g of urinary and reproductive tissues at day 7 post-phage treatment. (**D**) PFU recovered from vaginal swabs on day 1 through 6 of treatment. (**E**) Correlations of PFU and UTI89 Strep^R^ CFU recovered from vaginal swabs on day 1 through 6 of treatment. In a subset of mice, ΦHP3 or ΦCocktail was administered 30 min after UPEC inoculation and daily thereafter for an additional six days. (**F**) UTI89 Strep^R^ CFU recovered from vaginal swabs over seven days of treatment. (**G**) UTI89 Strep^R^ CFU/g of urinary and reproductive tissues at day 7 post-phage treatment. Experiments were performed in at least two independent replicate experiments. Points represent individual mice and lines represent medians with interquartile ranges (B-G). *n*=25-28 (A-C), *n*=28 (D-E), *n*=7-10 (F-G). Data were analyzed by Mantel-Cox test (A), Mann-Whitney *U* test with Benjamini, Krieger and Yekutieli correction for false discovery with a false discovery rate set at 5% (B), two-way ANOVA with Holm-Šídák’s multiple comparisons test (C), Friedman test with Dunn’s multiple comparisons test to day 1 values (D), Pearson correlation (E), or mixed-effects model with Dunnett’s multiple comparisons test (F-G), \**p*<0.05, \*\**p*<0.01, \*\*\**p*<0.001, \*\*\*\**p*<0.0001.

## DISCUSSION

While clinical evidence highlights the importance of UPEC vaginal colonization as a predictor of UTI risk(20, 22–24), with genetically identical vaginal and UTI isolates reported in some cases(74), there are limited options to control UPEC vaginal colonization. In this study, we explored the use of lytic bacteriophage as a UPEC urogenital decolonization strategy. We found that phage readily bound to human vaginal and bladder epithelial cells, reduced UPEC adherence in both cell types, and limited UPEC intracellular survival in vaginal cells *in vitro*. In a murine UPEC vaginal colonization model, although lytic phage was recovered and development of phage resistance was not observed, phage treatment only modestly reduced UPEC vaginal colonization, suggesting both a potential clinical benefit and a need for further optimization. In human vaginal samples, phage represent the largest portion (83%) of total viral sequences, are detected in up to 90% of human vaginal samples, and are predominantly predicted to be lysogenic (temperate) phage that incorporate into the bacterial genome rather than causing cell lysis(39, 40, 75). Endogenous phage predicted to infect *E. coli* are detected in about 10% of women(75) and are enriched in women with bacterial vaginosis(40), suggesting these lysogenic phages may confer a fitness advantage through enhanced virulence, stress response, or antibiotic resistance. While the therapeutic application of lytic phage in the vaginal tract is still in early stages, multiple studies have evaluated the potential of phage-derived endolysins to reduce vaginal pathobionts including group B *Streptococcus*(50) and *Gardnerella* spp.(76). Even so, there is substantial experimental work testing phage-mediated decolonization of other tissues(77) and across a range of model systems including invertebrate animals(78). A recent systematic review revealed 80% of experimental studies report phage-mediated reduction of the target organism at sites including the gastrointestinal tract, skin, lung, and urinary tract(77).

Building upon prior work, we selected ΦHP3, a *Myoviridae* family phage with broad efficacy against UPEC strains and established efficacy in human blood and urine(46, 53, 55, 63) to assess UPEC decolonization of the urogenital epithelium. Our four-phage cocktail included ΦHP3, as well as ΦHP3.1, to counter emerging bacterial resistors, and two phage, E17 and ES19, with superior anti-biofilm activity(54, 55). In contrast with other studies(72, 79), we did not consistently identify increased activity of the phage cocktail compared to single phage treatment either *in vitro* or *in vivo*. This could be explained by overlapping bacterial targets, believed to be primarily LPS for both HP3 and ES17(53, 54), or that the phage cocktail activity was primarily driven by ΦHP3, and that bacterial resistance and biofilm production were not key persistence mechanisms in our models.

Similar to prior work with an *Autographiviridae* phage(80), we observed phage-mediated reduction of UPEC interactions with HTB-9 cells and extended these studies to vaginal epithelial (VK2) cells. In both cell lines, reduction of UPEC was dependent on bacterial susceptibility to phage lysis since no effects of phage were observed using phage-resistant UTI89. Both ΦHP3 and ΦCocktail were somewhat more active on vaginal-derived cells compared to bladder-derived across adherence, invasion, and intracellular survival assays, even though lytic phage was recovered at similar levels from both lines, suggesting discordant phenotypes may be due to differences in extra- and intra-cellular localization of phage and bacteria between cell lines or impacts of cell culture media on phage activity. Similar to our findings, in a human bladder organoid model of non-transformed uroepithelial cells, ΦHP3 significantly reduced UTI89 adherence but did not impact established intracellular reservoirs(81). While eukaryotic cell uptake of phage may limit therapeutic efficacy of phage against extracellular bacteria(67), this uptake may allow phage to access intracellular bacterial populations. Phage reduction of intracellular bacterial reservoirs have been reported in murine macrophages(82) and human T24 bladder epithelial cells(83), and in this study we report phage-mediated reduction of intracellular UPEC in vaginal epithelial cells, but not HTB-9 bladder cells. A limitation is that we did not identify mechanism of phage entry in vaginal or bladder epithelial cells, although other phage have been described to undergo micropinocytosis, and clathrin- or caveolae-mediated endocytosis(84). Furthermore, we did not determine the eventual fate of the phage or phage DNA, although previous studies have shown that internalized phage are trafficked through the endolysosome and degraded(85). Furthermore, unlike prior studies with UTI89 and uroepithelial cells(86), we observed no differences in vaginal epithelial cell production of IL-8 in the presence of phage, suggesting augmentation of cellular immune responses did not contribute to our phenotypes.

Despite potent activity *in vitro*, phage have demonstrated mixed effects in animal models of pathogen colonization. Similar to our current study focused on UPEC urogenital colonization, prior studies have demonstrated that phage have limited efficacy in reducing exogenous or endogenous gut *E. coli*, even when active phage were recovered from the mucosal site(47, 87, 88). There are several plausible reasons for this. First, it is possible that the relative proportion, or MOI, of phage to bacterial host must be optimized to reduce bacterial colonization. Two phages used in our study, HP3 and ES17 (included in the cocktail), were effective in reducing extra-intestinal pathogenic *E. coli* murine gut colonization at a 10^10^ PFU dose, but not at a 10^9^ PFU dose(46). In contrast, a three-phage cocktail reduced UPEC gut colonization at both 10^5^ and 10^7^ doses(45), highlighting the range in activity of different phage and bacteria pairings. A second possibility is that the *in vivo* environment directly or indirectly impacts phage efficacy, with phage showing efficacy at in some tissues but not others(89). Phage binding of host mucins may impact *in vivo* efficacy, bringing them in closer proximity to the bacterial host such as the case for ΦES17 in the ΦCocktail(46, 90) or partially limiting the ability to bind bacterial hosts, such as is the case for ΦHP3(46). However, the similar activity between ΦHP3 and ΦCocktail suggest that mucin binding is not a strong driver of phenotypes in our models. Finally, the *in vivo* environment may provide the bacterial host a protected niche or low metabolic state that limits phage activity and replication. Similar to a study with *E. coli* mono-colonized mice(73), we observed positive correlations between recovered CFU and PFU with recovered bacteria post-phage treatment retaining phage susceptibility. Although we observed that endogenous murine *E. coli* vaginal isolates, a common member of the ^HMb^mice vaginal microbiota(51), were susceptible to the phage used in our experiments, we did not track endogenous *E. coli* levels before or after phage treatment. Thus, it is possible that endogenous *E. coli* was an additional variable influencing phage treatment success. As an additional limitation, we did not visualize the location of UPEC and phage *in vivo*. While all these possibilities may be at play in our model, neither reducing the UPEC inoculum nor altering the timing of phage treatment further augmented phage efficacy in reducing UPEC vaginal levels suggesting further optimization is needed.

Although beyond the scope of this work, it is possible that phage treatment may synergize with bacteria that outcompete UPEC in the urogenital environment. Combining phage with non-pathogenic *E. coli* reduces pathogenic *E. coli* colonization of the gastrointestinal tract *in vitro* (91) and *in vivo* (47); however, since *E. coli* is a lower abundance vaginal organism associated with a non-optimal vaginal microbiota(21), this is likely not a viable strategy for UPEC vaginal decolonization. Alternatively, inclusion of a vaginal *Lactobacillus* spp., associated with an optimal vaginal microbiota(92), that also displays anti-UPEC activity may serve as a promising approach to control UPEC vaginal colonization.

In summary, our study shows that phage can limit UPEC urogenital colonization across multiple experimental models; however, the biological factors dictating phage activity in this environment remain to be identified. Phage effectiveness required bacterial susceptibility to lysis. While spontaneous development of phage resistance was not readily detected in our models, experimentally acquired phage resistant strains displayed reduced colonization fitness *in vivo*. These findings deepen our knowledge of phage activity towards UPEC in the urogenital environment and support their continued development as a strategy to prevent UTIs.

## MATERIALS AND METHODS

### Bacterial strains and mammalian cell lines

Uropathogenic *E. coli* strain UTI89(93), an HP3 phage-resistant UTI89 derivative (UTI89-2)(53), here designated Φ^R^, a spontaneous streptomycin-resistant mutant of UTI89 (UTI89 Strep^R^)(51), UTI89 Φ^R^ Strep^R^ (this study using the same method), and UTI89-RFP Kan^R^(94). Strains were grown in Luria-Bertani (LB) broth overnight at 37°C with shaking and streptomycin (1000 µg/mL) or kanamycin (50 µg/mL) where applicable. *E. coli* DH5α (ATCC BAA-3219) was grown in LB broth to stationary phase and used as the bacterial host for phage plaque assays. Bacterial stocks were maintained at -80°C in 20% glycerol. Immortalized human vaginal epithelial cells (VK2/E6E7, ATCC CRL-2616) were grown in keratinocyte serum-free medium (KSFM) containing 50 mg/mL bovine pituitary extract and 0.1 ng/mL human recombinant epithelial growth factor. Human bladder epithelium carcinoma cells (5637, ATCC HTB-9) were grown in RPMI-1640 (Corning) containing 10% heat-inactivated fetal bovine serum. The cells were incubated at 37°C in 5% CO2 and 100% humidity and passaged every 3 to 5 days until achieving 80-90% confluence prior to seeding into 24-well tissue culture-treated plates.

### Bacteriophage strains, titering, and cocktail preparation

Bacteriophage HP3 (accession <u>KY608967</u>)(63), ES17 (accession <u>MN508615</u>), and ES19 (accession <u>MN508616</u>) were isolated from environmental sources (HP3) and wastewater (ES17 and ES19). HP3.1 is a derivative of HP3 (54). Individual phage stocks were prepared and purified by cesium chloride gradient centrifugation as described previously(63). Endotrap HD Columns (LIONEX GmbH) were used to deplete endotoxin per manufacturer’s instructions. Phage titers were determined by culture-based, double agar overlay plaque assays. Briefly, LB agar plates were overlaid with molten soft agar (LB containing 0.5% agarose), cooled to 4 °C, and inoculated with 10^7^ CFU of *E. coli* DH5α as prey. After solidification, serially diluted phage lysates were spotted on top agar, incubated overnight at 37°C, and plaques quantified to calculate phage titers in PFU/mL. Phage preps were stored in phage buffer(54) at 4°C until use.

### UPEC growth in the presence of phage

Overnight UPEC cultures were pelleted, resuspended in fresh LB or simulated vaginal fluid (SVF) medium at 1:10 dilution, and transferred to 96-well microtiter plates. SVF, a mimetic of human vaginal fluid, was prepared as described previously(65), with the exception that the final pH (5.0) was not further reduced to 4.5 with lactic acid. For phage challenge, 90 µL of adjusted bacterial inoculum in LB were mixed with 10 µL of ΦHP3 or Φ cocktail (10^9^ PFU/mL) in 96-well plates, performed in at least technical duplicate. Control wells were treated with LB or SVF media alone. Plates were incubated at 37°C with orbital shaking in a Tecan Infinite 200 plate reader and growth was monitored by measuring OD_600nm_ at 15 min intervals for 12 h. Area under the curve was calculated using GraphPad Prism v10.5.0

### Adherence and invasion assays

UPEC adhesion and invasion assays were performed as previously described(95) with some minor modifications. Confluent monolayers grown in 24-well tissue culture plates were replenished with fresh KSFM or RPMI 1640 + 10% FBS medium for VK2 and HTB-9 cells, respectively. For adherence assays, cells were pretreated with ՓHP3 or ΦCocktail (10^8^ PFU/well) for 1 h. Following phage treatment, selected wells were washed with PBS twice to retain only cell-adherent phage, while others remained unwashed to retain the total phage population within the well. Bacteria (UTI89 or UTI89 Φ^R^) were added at 10^6^ CFU/well, centrifuged for 2 min at 200×*g*, and incubated for 30 min at 37°C in 5% CO_2_. After 30 min of incubation, media was removed, and wells were washed at least three times with PBS to remove non-adherent extracellular bacteria. Cells were lifted by incubation with 100 µL of 0.025% Trypsin-EDTA for 8 min at 37°C. Thereafter, 400 µL of 0.025% Triton X-100 was added to each well. Cells were lysed with vigorous pipetting 30X and 10-fold serial dilutions of cell lysates were plated on LB agar to quantify CFU/well and percent adherence of the inoculum. For invasion assays, UPEC strains were incubated with host cell monolayers for 2 h, cells were washed with PBS three times, and then incubated with medium containing gentamicin (100 µg/mL) for 1 h to kill extracellular bacteria. Host cells were then trypsinized and permeabilized, serially diluted, and plated as described for adhesion assays to quantify invaded bacteria.

### Intracellular survival assays

As with adherence and invasion assays, VK2 and HTB-9 cell monolayers were seeded into 24-well tissue culture plates and replaced with fresh medium. Cells were infected with 10^6^ CFU/well UTI89 or UTI89 Φ^R^ (MOI of 10), plates were centrifuged at 200 × *g* for 2 min, and incubated at 37°C with 5% CO_2_ for UPEC internalization. After 2 h of incubation, cells were washed with PBS 1X and gentamicin was added at a concentration of 100 μg/mL for 1 h to kill extracellular bacteria. Infected cells were washed with PBS three times and treated with 100uL of 10^8^ PFU/well of ΦHP3 or ΦCocktail in fresh medium containing gentamicin (10 μg/mL) and incubated at 37°C. After 24 h of incubation with phage, cells were trypsinized and permeabilized, serially diluted, and plated as described above. Intracellular CFU were quantified the following day and expressed as CFU/well.

### Phage plaque assays

Phage plaques recovered from cell supernatant or cell lysate was determined using culture based double agar overlay. Briefly, supernatants and lysates containing phage were filtered (0.2 μm) and stored at 4°C until use. For the plaque assays, LB plates were overlaid with 3.5 mL of molten soft agarose (cooled to 45°C) containing 100 µl of stationary phase *E. coli* DH5α cultures. After solidification, samples were serially diluted and spotted (5 µl) on the “top agar” surface. Plates were incubated overnight at 37°C and plaques counted to calculate phage titers as PFU/mL.

### Cytokine assays

Confluent monolayers of VK2 cells were pretreated with ΦHP3 (10^8^ PFU/well) and incubated for 1h. Cell was challenged with 10^6^ CFU UTI89 (MOI of 10) and further incubated for 6 h at 37°C with 5% CO_2_. After incubation, cell supernatants were collected and stored at –20°C until analysis. Interleukin-8 (IL-8) in undiluted VK2 supernatant was quantified via ELISA per manufacturers’ instructions (R&D Systems, DY208-05).

### Fluorescence confocal microscopy

For confocal microscopy, VK2 and HTB-9 cells were seeded (80,000 cells/well) in 24-well plates (Cellvis, P-1.5H-N). The following day, cells were washed with PBS and replaced with 400 μL of fresh RPMI 1640 medium without phenol red. Monolayers were infected with 10^6^ CFU of UTI89-RFP and incubated for 3 h at 37°C with 5% CO_2_. Unadhered bacteria were washed with PBS and replaced with fresh RPMI 1640 media without phenol red. Purified phages were labelled with SYBR-Gold nucleic acid gel stain (2.5X concentration, Invitrogen, S11494) for 1 h in the dark at 4°C, followed by three washes with phage buffer on a Amicon-Ultra centrifugal unit 100-kDa membrane to remove excess stain. Washed phages were resuspended in a final volume of 1 mL in phage buffer at 10^9^ PFU/ mL. Labeled phage (10^9^ PFU/mL in 100 μL) were added to the preincubated wells containing cells and bacteria for 1 h. Following incubation, these wells were fixed with 4% PFA for 15 mins and washed with PBS. Cells were stained with Wheat Germ Agglutinin Alexa Fluor™ 647 Conjugate (WGA, 5µg/mL) and Hoechst 33342 nucleic acid stain (1 µg/mL) and incubated for 20 min. Cells were washed with PBS and replaced with fresh RPMI 1640 media. Both 2D and 3D images were captured using a Nikon Ti2 ECLIPSE confocal microscope with a 60 X oil-immersion objective. Z-stacks were acquired at a step size interval of 0.3 µm along the Z-axis. Time-lapse imaging was performed on VK2 and HTB-9 cells after adding labeled phage (10^9^ PFU/mL in 100 μL). The image acquisition was initiated immediately after phage addition with time lapse settings of 30 seconds intervals for a total of 10 min using a Nikon Ti2 ECLIPSE confocal microscope with a 60 X oil-immersion objective. All the images were processed in Fiji, and representative sequential frames (Substack) were extracted in ImageJ to visualize phage, UPEC, and host cell interactions.

### Animals

Animal experiments were approved by the Baylor College of Medicine (BCM) Institutional Animal Care and Use Committee (protocol AN-8233) and were performed under accepted veterinary standards. Mice were given food and water *ad libitum*. Humanized microbiota mice (^HMb^mice) were bred and maintained as described previously(52). Mice were randomly distributed so that each treatment group contained a similar age range (6-8 weeks). Mice were acclimatized for one week in the biohazard room prior to experiments.

### Murine UPEC vaginal colonization

Vaginal colonization studies were adapted from our previous studies (51, 96). Briefly, mice were synchronized with 0.5 mg β-estradiol administered intraperitoneally 24h prior to vaginal inoculation of 10^7^ CFU of UTI89 Strep^R^ or UTI89 Φ^R^ Strep^R^ in a 10 μL volume. Beginning 24h post-infection, mice received daily doses of purified UPEC-targeted ΦHP3, or ΦCocktail (10^8^ PFU in 10 μL) or phage buffer only as a mock control for 7 consecutive days. Vaginal swabs were collected daily just prior to phage treatment as described previously(51), and swabs were suspended in 100 μL of PBS. To quantify CFU, the swab sample (10 μL) was serially diluted and plated on LB agar containing streptomycin (1000 μg/mL). To quantify PFU, 20 µL of the remaining swab sample was mixed with an equal volume of chloroform, incubated for 15 min to kill bacteria, serially diluted in phage buffer, and spotted on to top agar plates (5 µL). After one week of phage treatment, urinary and reproductive organs of mice were harvested and homogenized as described previously(97), and plated on LB agar with streptomycin to assess UPEC dissemination. Several modifications were made to the protocol to test the impact on phage efficacy. In a subset of experiments, mice received 10^8^ PFU of phage administered intravaginally 30 minutes after UPEC infection (**Fig. 6F-G**). Additionally, mice were vaginally inoculated with lower dose of UTI89 Strep^R^ (10^5^ CFU) followed by administration of 10^8^ PFU of phage 24h post-infection.

### Statistics

*In vitro* experiments were performed at least three times independently with at least two technical duplicates. Mean values of independent experiments were used to represent experimental replicates for statistical analyses. *In vivo* experiments were conducted at least twice independently with individual mice serving as biological replicates. Experimental data were combined prior to statistical analyses. Two-way ANOVAs were used to compare bacterial growth curves, UTI89 adherence and invasion, intracellular survival (ΦHP3 treatment), VK2 IL-8 secretion, and murine tissue CFUs with multiple comparisons tests including uncorrected Fisher’s LSD or Holm-Šídák’s test as indicated in figure legends. A mixed-effects model with Dunnett’s multiple comparisons test was used for murine tissues with three group comparisons. One-way ANOVA with Holm-Šídák’s multiple comparisons test or Mann-Whitney *U* tests were used to compare area under the curves as indicated in figure legends. Kruskal-Wallis tests with Dunn’s multiple comparisons test were used to compare UTI89 Φ^R^ adherence and invasion. Paired *t* tests were used to compare intracellular survival (ΦCocktail treatment). Mann-Whitney *U* tests were used to compare adherent PFU, and vaginal swab CFUs with the addition of Benjamini, Krieger and Yekutieli correction for false discovery with a false discovery rate set at 5%. Mantel-Cox (log rank) tests were used to compare percent colonization over time. Pearson correlations were used to correlate CFU across tissues and CFU and PFU within the same tissue. Friedman test with Dunn’s multiple comparisons test were used to compare vaginal PFU to day 1 values to subsequent timepoints. Statistical analyses were performed using Prism, v10.5.0 (GraphPad Software Inc., La Jolla, CA). *P* values of <0.05 were considered statistically significant.

## AUTHOR CONTRIBUTIONS

BJ and KP conceived and designed experiments. BJ, JJZ, CS, ZAH, ABL, and DK performed experiments. BJ and KP analyzed and interpreted results. AT prepared phage stocks and RAB, AWM, and KP acquired funding. BJ and KP drafted the manuscript. All authors contributed the discussion and manuscript edits.

## ACKNOWLEDGEMENTS

We are grateful to the vivarium staff at BCM for animal husbandry, and Colleen Ardis for animal colony management. This research was supported by NIH U19 grant (AI157981) to KAP, RAB, and AWM. JJZ was supported by an NIH F31 training grant (DK136201). DK is supported by the Early Career Award Program grant from Thrasher Research Fund.

## Declaration of Interests Statement

AWM has equity in a biotech start-up, Phiogen. Baylor College of Medicine and AWM. have filed for intellectual property on therapeutic phages. IUM serves on the scientific advisory board of Seed Health.

## REFERENCES

1. Yang X, Chen H, Zheng Y, Qu S, Wang H, Yi F. 2022. Disease burden and long-term trends of urinary tract infections: A worldwide report. Front Public Health 10:888205.

2. Foxman B. 2014. Urinary tract infection syndromes: occurrence, recurrence, bacteriology, risk factors, and disease burden. Infect Dis Clin North Am 28:1–13.

3. Flores-Mireles AL, Walker JN, Caparon M, Hultgren SJ. 2015. Urinary tract infections: epidemiology, mechanisms of infection and treatment options. Nat Rev Microbiol 13:269–84.

4. Ronald A. 2002. The etiology of urinary tract infection: traditional and emerging pathogens. Am J Med 113 Suppl 1A:14S–19S.

5. Glover M, Moreira CG, Sperandio V, Zimmern P. 2014. Recurrent urinary tract infections in healthy and nonpregnant women. Urol Sci 25:1–8.

6. Russo TA, Stapleton A, Wenderoth S, Hooton TM, Stamm WE. 1995. Chromosomal restriction fragment length polymorphism analysis of Escherichia coli strains causing recurrent urinary tract infections in young women. J Infect Dis 172:440–5.

7. Koljalg S, Truusalu K, Vainumae I, Stsepetova J, Sepp E, Mikelsaar M. 2009. Persistence of Escherichia coli clones and phenotypic and genotypic antibiotic resistance in recurrent urinary tract infections in childhood. J Clin Microbiol 47:99–105.

8. Moreno E, Andreu A, Perez T, Sabate M, Johnson JR, Prats G. 2006. Relationship between Escherichia coli strains causing urinary tract infection in women and the dominant faecal flora of the same hosts. Epidemiol Infect 134:1015–23.

9. Chen SL, Wu M, Henderson JP, Hooton TM, Hibbing ME, Hultgren SJ, Gordon JI. 2013. Genomic diversity and fitness of E. coli strains recovered from the intestinal and urinary tracts of women with recurrent urinary tract infection. Sci Transl Med 5:184ra60.

10. Forde BM, Roberts LW, Phan MD, Peters KM, Fleming BA, Russell CW, Lenherr SM, Myers JB, Barker AP, Fisher MA, Chong TM, Yin WF, Chan KG, Schembri MA, Mulvey MA, Beatson SA. 2019. Population dynamics of an Escherichia coli ST131 lineage during recurrent urinary tract infection. Nat Commun 10:3643.

11. Tchesnokova VL, Rechkina E, Chan D, Haile HG, Larson L, Ferrier K, Schroeder DW, Solyanik T, Shibuya S, Hansen K, Ralston JD, Riddell K, Scholes D, Sokurenko EV. 2020. Pandemic Uropathogenic Fluoroquinolone-resistant Escherichia coli Have Enhanced Ability to Persist in the Gut and Cause Bacteriuria in Healthy Women. Clin Infect Dis 70:937–939.

12. Thanert R, Reske KA, Hink T, Wallace MA, Wang B, Schwartz DJ, Seiler S, Cass C, Burnham CA, Dubberke ER, Kwon JH, Dantas G. 2019. Comparative Genomics of Antibiotic-Resistant Uropathogens Implicates Three Routes for Recurrence of Urinary Tract Infections. mBio 10.

13. Worby CJ, Schreiber HLt, Straub TJ, van Dijk LR, Bronson RA, Olson BS, Pinkner JS, Obernuefemann CLP, Munoz VL, Paharik AE, Azimzadeh PN, Walker BJ, Desjardins CA, Chou WC, Bergeron K, Chapman SB, Klim A, Manson AL, Hannan TJ, Hooton TM, Kau AL, Lai HH, Dodson KW, Hultgren SJ, Earl AM. 2022. Longitudinal multi-omics analyses link gut microbiome dysbiosis with recurrent urinary tract infections in women. Nat Microbiol 7:630–639.

14. Neugent ML, Kumar A, Hulyalkar NV, Lutz KC, Nguyen VH, Fuentes JL, Zhang C, Nguyen A, Sharon BM, Kuprasertkul A, Arute AP, Ebrahimzadeh T, Natesan N, Xing C, Shulaev V, Li Q, Zimmern PE, Palmer KL, De Nisco NJ. 2022. Recurrent urinary tract infection and estrogen shape the taxonomic ecology and function of the postmenopausal urogenital microbiome. Cell Rep Med 3:100753.

15. Vaughan MH, Mao J, Karstens LA, Ma L, Amundsen CL, Schmader KE, Siddiqui NY. 2021. The Urinary Microbiome in Postmenopausal Women with Recurrent Urinary Tract Infections. J Urol 206:1222–1231.

16. Scalise ML, Garimano N, Sanz M, Padola NL, Leonino P, Pereyra A, Casale R, Amaral MM, Sacerdoti F, Ibarra C. 2022. Detection of Shiga Toxin-Producing Escherichia coli (STEC) in the Endocervix of Asymptomatic Pregnant Women. Can STEC Be a Risk Factor for Adverse Pregnancy Outcomes? Front Endocrinol (Lausanne) 13:945736.

17. Chow AW, Percival-Smith R, Bartlett KH, Goldring AM, Morrison BJ. 1986. Vaginal colonization with Escherichia coli in healthy women. Determination of relative risks by quantitative culture and multivariate statistical analysis. Am J Obstet Gynecol 154:120–6.

18. Gupta K, Stapleton AE, Hooton TM, Roberts PL, Fennell CL, Stamm WE. 1998. Inverse association of H2O2-producing lactobacilli and vaginal Escherichia coli colonization in women with recurrent urinary tract infections. J Infect Dis 178:446–50.

19. Hooton TM, Scholes D, Gupta K, Stapleton AE, Roberts PL, Stamm WE. 2005. Amoxicillin-clavulanate vs ciprofloxacin for the treatment of uncomplicated cystitis in women: a randomized trial. JAMA 293:949–55.

20. Hooton TM, Roberts PL, Stapleton AE. 2012. Cefpodoxime vs ciprofloxacin for short-course treatment of acute uncomplicated cystitis: a randomized trial. JAMA 307:583–9.

21. Boutouchent N, Vu TNA, Landraud L, Kennedy SP. 2024. Urogenital colonization and pathogenicity of E. Coli in the vaginal microbiota during pregnancy. Sci Rep 14:25523.

22. Pfau A, Sacks T. 1981. The bacterial flora of the vaginal vestibule, urethra and vagina in premenopausal women with recurrent urinary tract infections. J Urol 126:630–4.

23. Czaja CA, Stamm WE, Stapleton AE, Roberts PL, Hawn TR, Scholes D, Samadpour M, Hultgren SJ, Hooton TM. 2009. Prospective cohort study of microbial and inflammatory events immediately preceding Escherichia coli recurrent urinary tract infection in women. J Infect Dis 200:528–36.

24. Stamey TA, Sexton CC. 1975. The role of vaginal colonization with enterobacteriaceae in recurrent urinary infections. J Urol 113:214–7.

25. Navas-Nacher EL, Dardick F, Venegas MF, Anderson BE, Schaeffer AJ, Duncan JL. 2001. Relatedness of Escherichia coli colonizing women longitudinally. Mol Urol 5:31–6.

26. Stapleton AE. 2016. The Vaginal Microbiota and Urinary Tract Infection. Microbiol Spectr 4.

27. Dolk FCK, Pouwels KB, Smith DRM, Robotham JV, Smieszek T. 2018. Antibiotics in primary care in England: which antibiotics are prescribed and for which conditions? J Antimicrob Chemother 73:ii2-ii10.

28. Shah M, Barbosa TM, Stack G, Fleming A. 2024. Trends in antibiotic prescribing in primary care out-of-hours doctors’ services in Ireland. JAC Antimicrob Resist 6:dlae009.

29. Shively NR, Buehrle DJ, Clancy CJ, Decker BK. 2018. Prevalence of Inappropriate Antibiotic Prescribing in Primary Care Clinics within a Veterans Affairs Health Care System. Antimicrob Agents Chemother 62.

30. Aabenhus R, Hansen MP, Siersma V, Bjerrum L. 2017. Clinical indications for antibiotic use in Danish general practice: results from a nationwide electronic prescription database. Scand J Prim Health Care 35:162–169.

31. Kot B. 2019. Antibiotic Resistance Among Uropathogenic Escherichia coli. Pol J Microbiol 68:403–415.

32. Smith HS, Hughes JP, Hooton TM, Roberts P, Scholes D, Stergachis A, Stapleton A, Stamm WE. 1997. Antecedent antimicrobial use increases the risk of uncomplicated cystitis in young women. Clin Infect Dis 25:63–8.

33. Tenney J, Hudson N, Alnifaidy H, Li JTC, Fung KH. 2018. Risk factors for aquiring multidrug-resistant organisms in urinary tract infections: A systematic literature review. Saudi Pharm J 26:678–684.

34. Rosen DA, Hooton TM, Stamm WE, Humphrey PA, Hultgren SJ. 2007. Detection of intracellular bacterial communities in human urinary tract infection. PLoS Med 4:e329.

35. De Nisco NJ, Neugent M, Mull J, Chen L, Kuprasertkul A, de Souza Santos M, Palmer KL, Zimmern P, Orth K. 2019. Direct Detection of Tissue-Resident Bacteria and Chronic Inflammation in the Bladder Wall of Postmenopausal Women with Recurrent Urinary Tract Infection. J Mol Biol 431:4368–4379.

36. Brannon JR, Dunigan TL, Beebout CJ, Ross T, Wiebe MA, Reynolds WS, Hadjifrangiskou M. 2020. Invasion of vaginal epithelial cells by uropathogenic Escherichia coli. Nat Commun 11:2803.

37. Mysorekar IU, Hultgren SJ. 2006. Mechanisms of uropathogenic Escherichia coli persistence and eradication from the urinary tract. Proc Natl Acad Sci U S A 103:14170–5.

38. Liu Q, Zuo T, Lu W, Yeoh YK, Su Q, Xu Z, Tang W, Yang K, Zhang F, Lau LHS, Lui RNS, Chin ML, Wong R, Cheung CP, Zhu W, Chan PKS, Chan FKL, Lui GC, Ng SC. 2022. Longitudinal Evaluation of Gut Bacteriomes and Viromes after Fecal Microbiota Transplantation for Eradication of Carbapenem-Resistant Enterobacteriaceae. mSystems 7:e0151021.

39. Jakobsen RR, Haahr T, Humaidan P, Jensen JS, Kot WP, Castro-Mejia JL, Deng L, Leser TD, Nielsen DS. 2020. Characterization of the Vaginal DNA Virome in Health and Dysbiosis. Viruses 12.

40. Madere FS, Sohn M, Winbush AK, Barr B, Grier A, Palumbo C, Java J, Meiring T, Williamson AL, Bekker LG, Adler DH, Monaco CL. 2022. Transkingdom Analysis of the Female Reproductive Tract Reveals Bacteriophages form Communities. Viruses 14.

41. Hugerth LW, Krog MC, Vomstein K, Du J, Bashir Z, Kaldhusdal V, Fransson E, Engstrand L, Nielsen HS, Schuppe-Koistinen I. 2024. Defining Vaginal Community Dynamics: daily microbiome transitions, the role of menstruation, bacteriophages, and bacterial genes. Microbiome 12:153.

42. Zulk JJ, Patras KA, Maresso AW. 2024. The rise, fall, and resurgence of phage therapy for urinary tract infection. EcoSal Plus 12:eesp00292023.

43. Leitner L, Ujmajuridze A, Chanishvili N, Goderdzishvili M, Chkonia I, Rigvava S, Chkhotua A, Changashvili G, McCallin S, Schneider MP, Liechti MD, Mehnert U, Bachmann LM, Sybesma W, Kessler TM. 2021. Intravesical bacteriophages for treating urinary tract infections in patients undergoing transurethral resection of the prostate: a randomised, placebo-controlled, double-blind clinical trial. Lancet Infect Dis 21:427–436.

44. Kim P, Sanchez AM, Penke TJR, Tuson HH, Kime JC, McKee RW, Slone WL, Conley NR, McMillan LJ, Prybol CJ, Garofolo PM. 2024. Safety, pharmacokinetics, and pharmacodynamics of LBP-EC01, a CRISPR-Cas3-enhanced bacteriophage cocktail, in uncomplicated urinary tract infections due to Escherichia coli (ELIMINATE): the randomised, open-label, first part of a two-part phase 2 trial. Lancet Infect Dis 24:1319–1332.

45. Galtier M, De Sordi L, Maura D, Arachchi H, Volant S, Dillies MA, Debarbieux L. 2016. Bacteriophages to reduce gut carriage of antibiotic resistant uropathogens with low impact on microbiota composition. Environ Microbiol 18:2237–45.

46. Green SI, Gu Liu C, Yu X, Gibson S, Salmen W, Rajan A, Carter HE, Clark JR, Song X, Ramig RF, Trautner BW, Kaplan HB, Maresso AW. 2021. Targeting of Mammalian Glycans Enhances Phage Predation in the Gastrointestinal Tract. mBio 12.

47. Porter SB, Johnston BD, Kisiela D, Clabots C, Sokurenko EV, Johnson JR. 2022. Bacteriophage Cocktail and Microcin-Producing Probiotic Escherichia coli Protect Mice Against Gut Colonization With Multidrug-Resistant Escherichia coli Sequence Type 131. Front Microbiol 13:887799.

48. Landlinger C, Tisakova L, Oberbauer V, Schwebs T, Muhammad A, Latka A, Van Simaey L, Vaneechoutte M, Guschin A, Resch G, Swidsinski S, Swidsinski A, Corsini L. 2021. Engineered Phage Endolysin Eliminates Gardnerella Biofilm without Damaging Beneficial Bacteria in Bacterial Vaginosis Ex Vivo. Pathogens 10.

49. Castro J, Sousa LGV, Franca A, Podpera Tisakova L, Corsini L, Cerca N. 2022. Exploiting the Anti-Biofilm Effect of the Engineered Phage Endolysin PM-477 to Disrupt In Vitro Single- and Dual-Species Biofilms of Vaginal Pathogens Associated with Bacterial Vaginosis. Antibiotics (Basel) 11.

50. Cheng Q, Nelson D, Zhu S, Fischetti VA. 2005. Removal of group B streptococci colonizing the vagina and oropharynx of mice with a bacteriophage lytic enzyme. Antimicrob Agents Chemother 49:111–7.

51. Mejia ME, Mercado-Evans V, Zulk JJ, Ottinger S, Ruiz K, Ballard MB, Fowler SW, Britton RA, Patras KA. 2023. Vaginal microbial dynamics and pathogen colonization in a humanized microbiota mouse model. NPJ Biofilms Microbiomes 9:87.

52. Collins J, Auchtung JM, Schaefer L, Eaton KA, Britton RA. 2015. Humanized microbiota mice as a model of recurrent Clostridium difficile disease. Microbiome 3:35.

53. Zulk JJ, Clark JR, Ottinger S, Ballard MB, Mejia ME, Mercado-Evans V, Heckmann ER, Sanchez BC, Trautner BW, Maresso AW, Patras KA. 2022. Phage Resistance Accompanies Reduced Fitness of Uropathogenic Escherichia coli in the Urinary Environment. mSphere 7:e0034522.

54. Salazar KC, Ma L, Green SI, Zulk JJ, Trautner BW, Ramig RF, Clark JR, Terwilliger AL, Maresso AW. 2021. Antiviral Resistance and Phage Counter Adaptation to Antibiotic-Resistant Extraintestinal Pathogenic Escherichia coli. mBio 12.

55. Sanchez BC, Heckmann ER, Green SI, Clark JR, Kaplan HB, Ramig RF, Hines-Munson C, Skelton F, Trautner BW, Maresso AW. 2022. Development of Phage Cocktails to Treat E. coli Catheter-Associated Urinary Tract Infection and Associated Biofilms. Front Microbiol 13:796132.

56. Ma L, Green SI, Trautner BW, Ramig RF, Maresso AW. 2018. Metals Enhance the Killing of Bacteria by Bacteriophage in Human Blood. Sci Rep 8:2326.

57. Moses S, Vagima Y, Tidhar A, Aftalion M, Mamroud E, Rotem S, Steinberger-Levy I. 2021. Characterization of Yersinia pestis Phage Lytic Activity in Human Whole Blood for the Selection of Efficient Therapeutic Phages. Viruses 13.

58. Mutti M, Moreno DS, Restrepo-Cordoba M, Visram Z, Resch G, Corsini L. 2023. Phage activity against Staphylococcus aureus is impaired in plasma and synovial fluid. Sci Rep 13:18204.

59. Dalmasso M, de Haas E, Neve H, Strain R, Cousin FJ, Stockdale SR, Ross RP, Hill C. 2015. Isolation of a Novel Phage with Activity against Streptococcus mutans Biofilms. PLoS One 10:e0138651.

60. Pavlova SI, Kilic AO, Mou SM, Tao L. 1997. Phage infection in vaginal lactobacilli: an in vitro study. Infect Dis Obstet Gynecol 5:36–44.

61. Kilic AO, Pavlova SI, Alpay S, Kilic SS, Tao L. 2001. Comparative study of vaginal Lactobacillus phages isolated from women in the United States and Turkey: prevalence, morphology, host range, and DNA homology. Clin Diagn Lab Immunol 8:31–9.

62. Damelin LH, Paximadis M, Mavri-Damelin D, Birkhead M, Lewis DA, Tiemessen CT. 2011. Identification of predominant culturable vaginal Lactobacillus species and associated bacteriophages from women with and without vaginal discharge syndrome in South Africa. J Med Microbiol 60:180–183.

63. Green SI, Kaelber JT, Ma L, Trautner BW, Ramig RF, Maresso AW. 2017. Bacteriophages from ExPEC Reservoirs Kill Pandemic Multidrug-Resistant Strains of Clonal Group ST131 in Animal Models of Bacteremia. Sci Rep 7:46151.

64. Terwilliger A, Clark J, Karris M, Hernandez-Santos H, Green S, Aslam S, Maresso A. 2021. Phage Therapy Related Microbial Succession Associated with Successful Clinical Outcome for a Recurrent Urinary Tract Infection. Viruses 13.

65. Brandt K, Barrangou R. 2020. Adaptive response to iterative passages of five Lactobacillus species in simulated vaginal fluid. BMC Microbiol 20:339.

66. Shan J, Ramachandran A, Thanki AM, Vukusic FBI, Barylski J, Clokie MRJ. 2018. Bacteriophages are more virulent to bacteria with human cells than they are in bacterial culture; insights from HT-29 cells. Sci Rep 8:5091.

67. Bichet MC, Chin WH, Richards W, Lin YW, Avellaneda-Franco L, Hernandez CA, Oddo A, Chernyavskiy O, Hilsenstein V, Neild A, Li J, Voelcker NH, Patwa R, Barr JJ. 2021. Bacteriophage uptake by mammalian cell layers represents a potential sink that may impact phage therapy. iScience 24:102287.

68. Nale JY, Ahmed B, Haigh R, Shan J, Phothaworn P, Thiennimitr P, Garcia A, AbuOun M, Anjum MF, Korbsrisate S, Galyov EE, Malik DJ, Clokie MRJ. 2023. Activity of a Bacteriophage Cocktail to Control Salmonella Growth Ex Vivo in Avian, Porcine, and Human Epithelial Cell Cultures. Phage (New Rochelle) 4:11–25.

69. Mosier-Boss PA, Lieberman SH, Andrews JM, Rohwer FL, Wegley LE, Breitbart M. 2003. Use of fluorescently labeled phage in the detection and identification of bacterial species. Appl Spectrosc 57:1138–44.

70. Robinson CK, Saenkham-Huntsinger P, Hanson BS, Adams LG, Subashchandrabose S. 2022. Vaginal Inoculation of Uropathogenic Escherichia coli during Estrus Leads to Genital and Renal Colonization. Infect Immun 90:e0053221.

71. Schlievert PM, Kilgore SH, Benavides A, Klingelhutz AJ. 2022. Pathogen Stimulation of Interleukin-8 from Human Vaginal Epithelial Cells through CD40. Microbiol Spectr 10:e0010622.

72. Buttimer C, Sutton T, Colom J, Murray E, Bettio PH, Smith L, Bolocan AS, Shkoporov A, Oka A, Liu B, Herzog JW, Sartor RB, Draper LA, Ross RP, Hill C. 2022. Impact of a phage cocktail targeting Escherichia coli and Enterococcus faecalis as members of a gut bacterial consortium in vitro and in vivo. Front Microbiol 13:936083.

73. Weiss M, Denou E, Bruttin A, Serra-Moreno R, Dillmann ML, Brussow H. 2009. In vivo replication of T4 and T7 bacteriophages in germ-free mice colonized with Escherichia coli. Virology 393:16–23.

74. Sekito T, Sadahira T, Hirakawa H, Ishii A, Wada K, Araki M. 2024. Homology of Escherichia coli isolated from urine and vagina and their antimicrobial susceptibility in postmenopausal women with recurrent cystitis. J Infect Chemother 30:1081–1084.

75. da Costa AC, Moron AF, Forney LJ, Linhares IM, Sabino E, Costa SF, Mendes-Correa MC, Witkin SS. 2021. Identification of bacteriophages in the vagina of pregnant women: a descriptive study. BJOG 128:976–982.

76. Arroyo-Moreno S, Cummings M, Corcoran DB, Coffey A, McCarthy RR. 2022. Identification and characterization of novel endolysins targeting Gardnerella vaginalis biofilms to treat bacterial vaginosis. NPJ Biofilms Microbiomes 8:29.

77. Fang Q, Yin X, He Y, Feng Y, Zhang L, Luo H, Yin G, McNally A, Zong Z. 2024. Safety and efficacy of phage application in bacterial decolonisation: a systematic review. Lancet Microbe 5:e489–e499.

78. Eddoubaji Y, Aldeia C, Campos-Madueno EI, Moser AI, Kundlacz C, Perreten V, Hilty M, Endimiani A. 2024. A new in vivo model of intestinal colonization using Zophobas morio larvae: testing hyperepidemic ESBL- and carbapenemase-producing Escherichia coli clones. Front Microbiol 15:1381051.

79. Kim MK, Chen Q, Echterhof A, Pennetzdorfer N, McBride RC, Banaei N, Burgener EB, Milla CE, Bollyky PL. 2024. A blueprint for broadly effective bacteriophage-antibiotic cocktails against bacterial infections. Nat Commun 15:9987.

80. Li X, Xu S, Tan L, Yan X, Wang X, Li Z, Chen L, Zhang W. 2025. Characterization of a novel phage vB_EcoP_P64441 and its potential role in controlling uropathogenic Escherichia coli (UPEC) and biofilms formation. Virology 609:110570.

81. Zulk JJ, Robertson CM, Ottinger S, Kambal A, Tostado AR, Fleck RC, Shea AE, Coarfa C, Blutt SE, Maresso AW, Patras KA. 2025. Human bladder organoids model urinary tract infection and bacteriophage therapy. bioRxiv doi:10.1101/2025.07.30.667685.

82. Capparelli R, Parlato M, Borriello G, Salvatore P, Iannelli D. 2007. Experimental phage therapy against Staphylococcus aureus in mice. Antimicrob Agents Chemother 51:2765–73.

83. Moller-Olsen C, Ho SFS, Shukla RD, Feher T, Sagona AP. 2018. Engineered K1F bacteriophages kill intracellular Escherichia coli K1 in human epithelial cells. Sci Rep 8:17559.

84. Goswami A, Sharma PR, Agarwal R. 2021. Combatting intracellular pathogens using bacteriophage delivery. Crit Rev Microbiol 47:461–478.

85. Lehti TA, Pajunen MI, Skog MS, Finne J. 2017. Internalization of a polysialic acid-binding Escherichia coli bacteriophage into eukaryotic neuroblastoma cells. Nat Commun 8:1915.

86. Kongsomboonchoke P, Mongkolkarvin P, Khunti P, Vijitphichiankul J, Nonejuie P, Thiennimitr P, Chaikeeratisak V. 2025. Rapid formulation of a genetically diverse phage cocktail targeting uropathogenic Escherichia coli infections using the UTI89 model. Sci Rep 15:12832.

87. Javaudin F, Bemer P, Batard E, Montassier E. 2021. Impact of Phage Therapy on Multidrug-Resistant Escherichia coli Intestinal Carriage in a Murine Model. Microorganisms 9.

88. Chibani-Chennoufi S, Sidoti J, Bruttin A, Kutter E, Sarker S, Brussow H. 2004. In vitro and in vivo bacteriolytic activities of Escherichia coli phages: implications for phage therapy. Antimicrob Agents Chemother 48:2558–69.

89. Dufour N, Clermont O, La Combe B, Messika J, Dion S, Khanna V, Denamur E, Ricard JD, Debarbieux L, ColoColi g. 2016. Bacteriophage LM33_P1, a fast-acting weapon against the pandemic ST131-O25b:H4 Escherichia coli clonal complex. J Antimicrob Chemother 71:3072–3080.

90. Barr JJ, Auro R, Furlan M, Whiteson KL, Erb ML, Pogliano J, Stotland A, Wolkowicz R, Cutting AS, Doran KS, Salamon P, Youle M, Rohwer F. 2013. Bacteriophage adhering to mucus provide a non-host-derived immunity. Proc Natl Acad Sci U S A 110:10771–6.

91. Forsyth JH, Barron NL, Scott L, Watson BNJ, Chisnall MAW, Meaden S, van Houte S, Raymond B. 2023. Decolonizing drug-resistant E. coli with phage and probiotics: breaking the frequency-dependent dominance of residents. Microbiology (Reading) 169.

92. Ottinger S, Robertson CM, Branthoover H, Patras KA. 2024. The human vaginal microbiota: from clinical medicine to models to mechanisms. Curr Opin Microbiol 77:102422.

93. Mulvey MA, Schilling JD, Hultgren SJ. 2001. Establishment of a persistent Escherichia coli reservoir during the acute phase of a bladder infection. Infect Immun 69:4572–9.

94. Mora-Bau G, Platt AM, van Rooijen N, Randolph GJ, Albert ML, Ingersoll MA. 2015. Macrophages Subvert Adaptive Immunity to Urinary Tract Infection. PLoS Pathog 11:e1005044.

95. Zulk JJ, Clark JR, Ottinger S, Ballard MB, Mejia ME, Mercado-Evans V, Heckmann ER, Sanchez BC, Trautner BW, Maresso AW. 2022. Phage resistance accompanies reduced fitness of uropathogenic Escherichia coli in the urinary environment. Msphere 7:e00345–22.

96. Patras KA, Doran KS. 2016. A murine model of group B Streptococcus vaginal colonization. JoVE (Journal of Visualized Experiments):e54708.

97. Mercado-Evans V, Mejia ME, Zulk JJ, Ottinger S, Hameed ZA, Serchejian C, Marunde MG, Robertson CM, Ballard MB, Ruano SH, Korotkova N, Flores AR, Pennington KA, Patras KA. 2024. Gestational diabetes augments group B Streptococcus infection by disrupting maternal immunity and the vaginal microbiota. Nat Commun 15:1035.

